# An inverse Lansing effect: Older mothers produce offspring with improved development and growth efficiency

**DOI:** 10.1101/2025.10.16.682747

**Authors:** Keshi Zhang, Peter Schausberger, Greg Holwell, Zhi-Qiang Zhang

## Abstract

The Lansing effect predicts a decline in offspring performance with increasing maternal age. Maternal age and diet can influence offspring development and fitness via maternal effects, but how these two factors interact remains poorly understood. We examined how maternal age at oviposition and dietary conditions affect offspring developmental plasticity in a thelytokous predatory mite (*Amblyseius herbicolus*). Mothers were provided either a restricted or abundant prey diet, and their offspring were exposed to varying prey availability and monitored for hatching success, survival to adulthood, developmental time, size at maturity, and prey consumption. We addressed two main questions: How does maternal age affect offspring developmental time and size at maturity and does maternal diet modify the effect of maternal age on offspring? Our results suggest an inverse Lansing effect. Offspring of older mothers showed increased survival and reduced prey consumption without any compromise in terms of size at maturity. Interactions were found between maternal diet and age on offspring prey consumption and developmental plasticity. Notably, offspring from older, diet-restricted mothers achieved the best overall performance during development. Our study demonstrates that maternal age and diet jointly shape offspring development, and highlights the importance of incorporating maternal age into studies of maternal effects and phenotypic plasticity.

## 1 Introduction

To cope with environmental variability, organisms have evolved diverse strategies to optimise survival and reproduction (West-Eberhard, 1989). One such strategy is phenotypic plasticity, which is defined as the ability of a single genotype to produce different phenotypes in response to environmental variation (West-Eberhard, 2003). Phenotypic adjustments can occur during an organism’s development, a process known as developmental plasticity, which often results in irreversible trait changes (Piersma & Drent, 2003; West-Eberhard, 2005; Blanckenhorn, 2009; Taborsky, 2017). Developmental plasticity allows individuals to adjust life-history traits such as growth rate, development time, and size at maturity, particularly under suboptimal conditions such as food limitation or temperature stress (Stearns & Koella, 1986; Blanckenhorn, 1999; Bize et al., 2003). This ability is especially important in species with determinate growth, where development ceases upon maturity (Bize et al., 2003).

However, plastic responses are not confined to an individual’s own generation (Räsänen & Kruuk, 2007; Bonduriansky & Head, 2007). Increasing evidence shows that parental environmental experiences can shape offspring phenotypes via maternal and/or paternal effects, a form of non-genetic inheritance that is widespread across taxa (Bonduriansky & Day, 2009; Bonduriansky, 2012; Luquet & Tariel, 2016; Kuijper & Johnstone, 2018; Bell & Hellmann, 2019). Through mechanisms such as epigenetic modification, hormonal signalling, or differential provisioning, parents can influence offspring traits including body size, development, stress tolerance, and survival (Bonduriansky & Day, 2009; Bonduriansky, 2012; Luquet & Tariel, 2016; Kuijper & Johnstone, 2018; Bell & Hellmann, 2019; Groothuis et al., 2020). Parental effects can also alter offspring developmental trajectories (Plaistow et al., 2004; Luquet & Tariel, 2016; Groothuis et al., 2020).

Among intrinsic parental factors, maternal age has emerged as a key determinant of offspring phenotype (Plaistow et al., 2015; Perez et al., 2017; Bonduriansky et al., 2018; Hernández et al., 2020). The decline in offspring performance with maternal age—known as the Lansing effect or maternal effect senescence—has been found across a wide range of taxa, from rotifers and insects to birds and mammals (Lansing, 1947; Fox et al., 2003; Bouwhuis et al., 2015; Schroeder et al., 2015; Moorad & Nussey, 2016; Reichert et al., 2020; Krug et al., 2020; Ivimey-Cook et al., 2023). Maternal age effects can manifest as reduced hatching success, lower stress resistance, smaller size, or shortened lifespan in offspring (Halle et al., 2015; Bloch Qazi et al., 2017; Hernández et al., 2020). The adverse influences of advanced maternal age are often context-dependent and shaped by environmental conditions such as diet (Plaistow & Benton, 2009; Gribble et al., 2014; van Daalen et al., 2022).

Two main theoretical frameworks have been put forward to explain maternal age effects. The senescent parent hypothesis proposes that physiological deterioration reduces gamete quality or maternal provisioning, thereby reducing offspring fitness (Kong et al., 2012; Plaistow et al., 2015; Bloch Qazi et al., 2017). For example, maternal age is linked to oxidative damage in *Drosophila* oocytes (Fredriksson et al., 2012) and reduced nutrient provisioning in eggs of the wasp *Eupelmus vuilletti* (Muller et al. 2017). Such findings challenge the traditional separation between germline and soma, suggesting that age-related decline can penetrate the Weismann barrier (Monaghan & Metcalfe, 2019). By contrast, adaptive hypotheses predict age-related shifts in reproductive strategy (Goos et al. 2019). Both the asset protection principle and the reproductive restraint hypothesis suggest that older individuals may reduce reproductive effort and increase investment in somatic maintenance (Clark, 1994; Jehan et al., 2021). Conversely, the terminal investment hypothesis suggests increased reproductive effort late in life (Monaghan et al., 2020; Jehan et al., 2021). For instance, young soil mite (*Sancassania berlesei*) mothers produce more but smaller eggs, whereas older mothers produce fewer but larger ones (Plaistow et al., 2007).

Crucially, the strength and direction of maternal age effects often depend on environmental conditions, especially diet (Rollinson & Hutchings, 2013; van Daalen et al., 2022). Diet stress can exacerbate negative effects on offspring (Vijendravarma et al., 2010; Hafer et al., 2011), whereas mild diet restriction (i.e., caloric restriction) may extend maternal lifespan and sometimes even enhance offspring performance (Gribble et al., 2014; Hibshman et al., 2016). Interactions between maternal age and diet are predicted to strongly influence offspring developmental plasticity (van den Heuvel et al., 2016).

Despite growing recognition of these interactions, empirical studies remain scarce. To address this, we investigated the combined effects of maternal age and diet on the offspring of a thelytokous predatory mite, *Amblyseius herbicolus* (Chant) (Acari: Phytoseiidae). This asexual, oviparous species provides a system that eliminates confounding factors such as paternal genetic input and maternal care, enabling a clear focus on physiological and epigenetic maternal effects (Castonguay & Angers, 2012; Verhoeven & Preite, 2014; Groothuis et al., 2020; Vogt, 2021). Although no Lansing effect was previously detected in *A. herbicolus* when fed ad libitum prey, older females consistently produced smaller eggs, suggesting age-related shifts in provisioning (Zhang et al., 2024). How these shifts influence offspring developmental plasticity remains unclear.

Here, we tested two hypotheses:

1. Offspring of older mothers exhibit reduced developmental plasticity, expressed as narrower ranges of developmental time and body size at maturity.
2. Dietary restriction modulates maternal age effects, strengthening negative age effects under low-resource conditions.

By examining the interaction between maternal condition and offspring developmental plasticity, this study provides insights into our understanding of transgenerational effects and their ecological and evolutionary significance, especially in asexually reproducing species.

## 2 Materials and Methods

### 2.1 Study animals and feeding conditions

*Amblyseius herbicolus* is a generalist predatory mite, typically less than 500 μm in body length (McMurtry et al., 2013; Zhang & Zhang, 2021). It has five life stages (egg, larva, protonymph, deutonymph, and adult; FIGURE S1), a developmental period of about 1 week, and a lifespan of about 1 month under laboratory conditions (Liu et al., 2024a). We used a laboratory population of *A. herbicolus* to examine the influence of maternal age and diet on offspring developmental plasticity. The founding population (>30 adult females) was collected from avocado (*Persea americana* Mill.) leaves in an orchard in Te Puna, Tauranga, New Zealand. Species identification was confirmed based on morphological characteristics described by Ma et al. (2024). After collection, predatory mites were fed the dried fruit mite *Carpoglyphus lactis* L. (Acari: Carpoglyphidae), sourced from Bioforce Limited (Karaka, Auckland, New Zealand), for 3 months before the experiment (approximately six generations). Colonies were maintained on water-filled plates (Figure S2A; Zhang & Zhang, 2021) within plexiglass cabinets at 25 °C ± 1 °C, 80% ± 5% relative humidity, and a 16:8 h (light:dark) photoperiod.

### 2.2 Preparing prey eggs

Frozen *C. lactis* eggs were used as prey. These eggs are suitable food for predatory mites (Liu et al., 2024a). Using non-viable eggs reduced contamination from yeast-based rearing and the need for frequent cell changes. Eggs of *C. lactis* were collected following Liu et al. (2024b), frozen at −18 °C for about a week, thawed at room temperature for 20 min, and used within 1 month of freezing.

### 2.3 Experimental procedures

#### 2.3.1 Maternal generation

Approximately 50 *A. herbicolus* eggs (<16 h old) were transferred to new plates (FIGURE S2A) and reared with ad libitum access to mixed-stage *C. lactis*. Eggs were collected by placing 1 cm sewing threads in colonies overnight, producing synchronous cohorts. Development to adulthood requires ∼10 days, so new threads were placed on days 10–15 to collect newly laid eggs (<16 h old). Threads were replaced daily, and collected eggs were used in experiments (FIGURE 1A).

**FIGURE 1.**
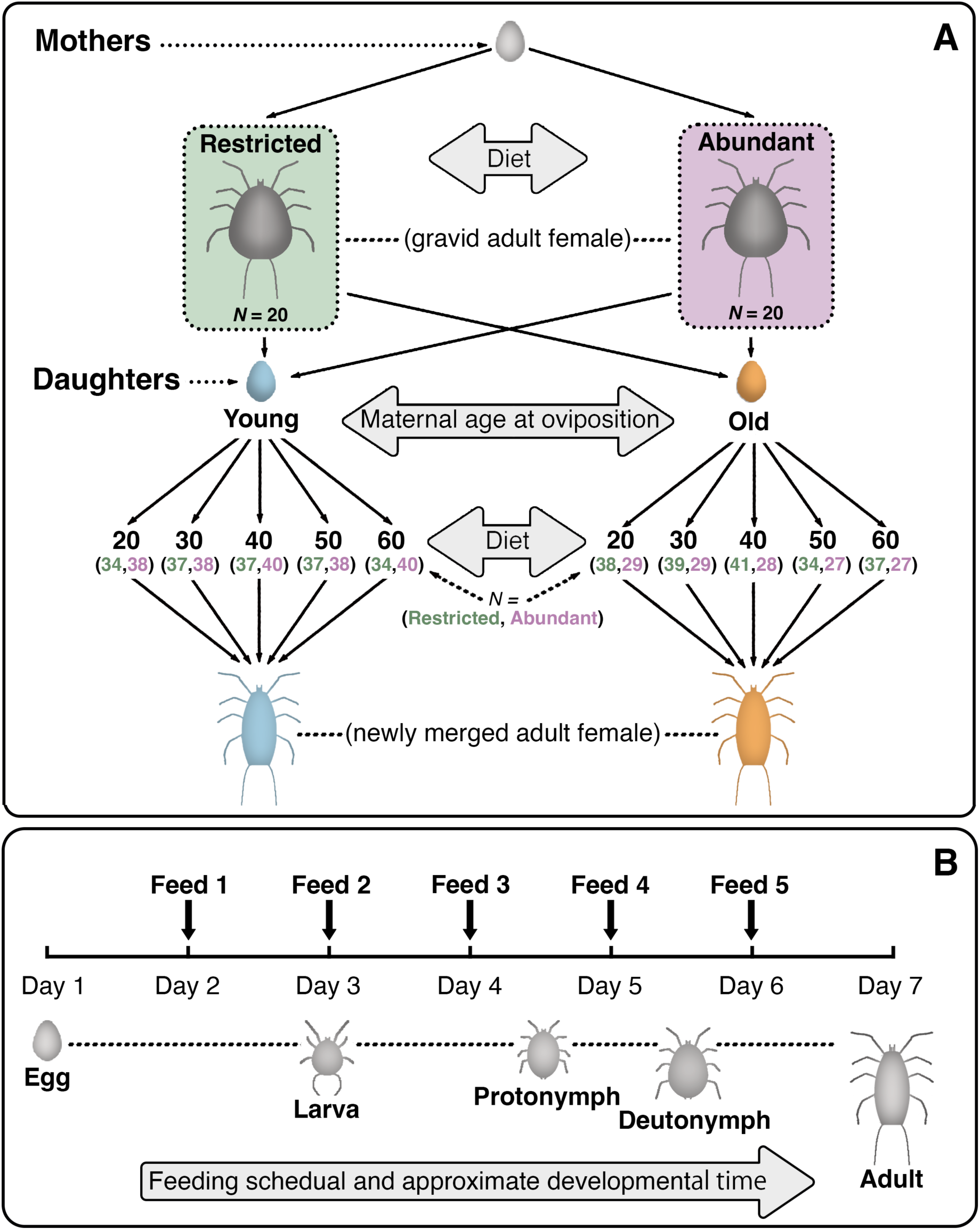
Schematic representation of the experimental design used in this study. A: Experimental setup for examining the effects of maternal age and dietary conditions on the developmental plasticity of *Amblyseius herbicolus* offspring. Sample sizes (*N*) are provided for each treatment group; for daughters, sample sizes are shown in brackets, with values for restricted and abundant maternal diets listed on the left and right, respectively. B: Feeding schedule (feeds 1–5) during the immature development of *A. herbicolus* for both mothers and daughters. Since the hatching time of *A. herbicolus* eggs exceeded 24 h, the first feeding occurred on day 2 to ensure newly hatched larvae could access food. To prevent an accumulation of uneaten prey before larval emergence, the second feeding was administered only after larvae were observed, typically on day 3. Subsequent feedings continued daily until the final prey provision.

These synchronous cohorts of *A. herbicolus* eggs were randomly assigned to one of two dietary treatments and reared individually until oviposition (FIGURE S2B), as follows:

1. Restricted diet: 30 prey eggs in total during immature development (6 per day for 5 days; FIGURE 1B), and 15 eggs per day during adulthood.
2. Abundant diet: 50 prey eggs in total during immature development (10 per day for 5 days; FIGURE 1B), and 40 eggs per day during adulthood.

Twenty eggs were allocated to each treatment group as replicates. Reproduction is thelytokous so mating was not required. Upon maturity, females were transferred to new rearing cells, and prey was replenished daily to facilitate oviposition.

#### 2.3.2 Offspring generation

Eggs (<16 h old) laid by mothers were assigned to two measurements:

1. Egg size measurement: egg volume (*V*) was estimated from length (*L*) and maximum breadth (*B*) using Narushin’s (2005) formula: *V* = (0.6057 − 0.0018*B*)*LB*^2^
2. Developmental plasticity: daughters were reared individually (FIGURE 1A) and provided with one of five prey densities during immature development: 4, 6, 8, 10, or 12 eggs per day for 5 days (20–60 eggs total; FIGURE 1B). Offspring were classified by maternal age at oviposition as ‘young’ or ‘old,’ with the threshold defined as half the mean oviposition period of mothers in each dietary treatment group.

### 2.4 Data collection

Both the mothers and daughters were monitored twice daily (08:00 and 17:30). For mothers, developmental time (egg to adult), prey consumption (number of prey eggs consumed), size at maturity (dorsal plate length), fecundity (lifetime oviposition), and lifespan (egg to death) were recorded. For offspring, hatching success, survival to adulthood, developmental time, prey consumption, and size at maturity were recorded. Adult size for both generations was determined by mounting mature individuals in Hoyer’s medium and examining them under a compound microscope (Walter & Krantz 2009).

### 2.5 Statistical analysis

Analyses were conducted in R (R Core Team, 2024) in RStudio (version 2024.09.1). Graphics and models were generated using *ggplot2* and *lme4* (Bates et al., 2015; Wickham, 2016). For the maternal generation, two eggs from the abundant group did not hatch, and one individual from the abundant group and two from the restricted group were lost during feeding. The remaining individuals survived to oviposition and were analysed. Maternal life-history traits were summarised as means ± standard errors of the mean (SEMs). Generalised linear models (GLMs) with Poisson or quasi-Poisson distributions tested the effect of dietary treatments, with model dispersion verified post hoc. Spearman’s correlations examined the relationships between maternal age, prey consumption, and daily oviposition. Egg volume was analysed with linear mixed-effects models (LMMs), including maternal diet and age as fixed effects and mother as a random effect.

For the offspring generation, logistic regressions tested the effects of maternal diet, maternal age, and offspring prey availability on hatching and survival (binomial response). Developmental time and size at maturity were analysed using LMMs with fixed factors (maternal diet, maternal age, and prey availability), a covariate (prey consumption), and mother as a random factor. Likelihood ratio tests (LRTs) were used to assess model fit. Preliminary models including prey availability showed no significant effect on developmental time (LRT: χ^2^ = 4.258, *df* = 4, *P* = 0.372) and size at maturity (χ^2^ = 6.174, *df* = 4, *P* = 0.187), and prey availability was therefore excluded from the final models. Pearson’s correlations assessed the relationships between prey consumption and developmental traits. Offspring prey consumption was analysed using a Poisson GLM, with maternal diet and age as fixed factors and prey consumption as a covariate. Statistical significance was set at α = 0.05.

## 3 RESULTS

### 3.1 Diet-induced influences on maternal parameters

Several life-history traits of mothers were significantly influenced by dietary conditions (TABLE S1). Individuals in the restricted diet group exhibited longer developmental and oviposition periods and an extended lifespan, whereas those in the abundant diet group showed higher daily oviposition rates, greater maximum daily fecundity, and larger body size. The post-oviposition period and lifetime fecundity did not differ between treatments, while the pre-oviposition period was marginally non-significant (TABLE S1).

Individuals given more prey consumed more during both immature development and adulthood (TABLE S1). After reaching adulthood, daily prey consumption declined with age in both dietary groups (Figure S3). Average daily oviposition ranged from one to two eggs in both treatment groups; in the restricted group, this rate remained consistent throughout the oviposition period, whereas in the abundant group it declined with maternal age (FIGURE S4).

Egg size was significantly influenced by dietary condition (LMM: Wald χ^2^ = 57.673, *df* = 1, *P* < 0.001) and maternal age at oviposition (Wald χ^2^ = 74.691, *df* = 1, *P* < 0.001). Mothers on the abundant diet produced larger eggs, and egg size declined with age. A significant interaction between diet and maternal age (Wald χ^2^ = 47.163, *df* = 1, *P* < 0.001) indicated that egg size decreased more sharply with age in mothers on the restricted diet (FIGURE 2).

**FIGURE 2.**
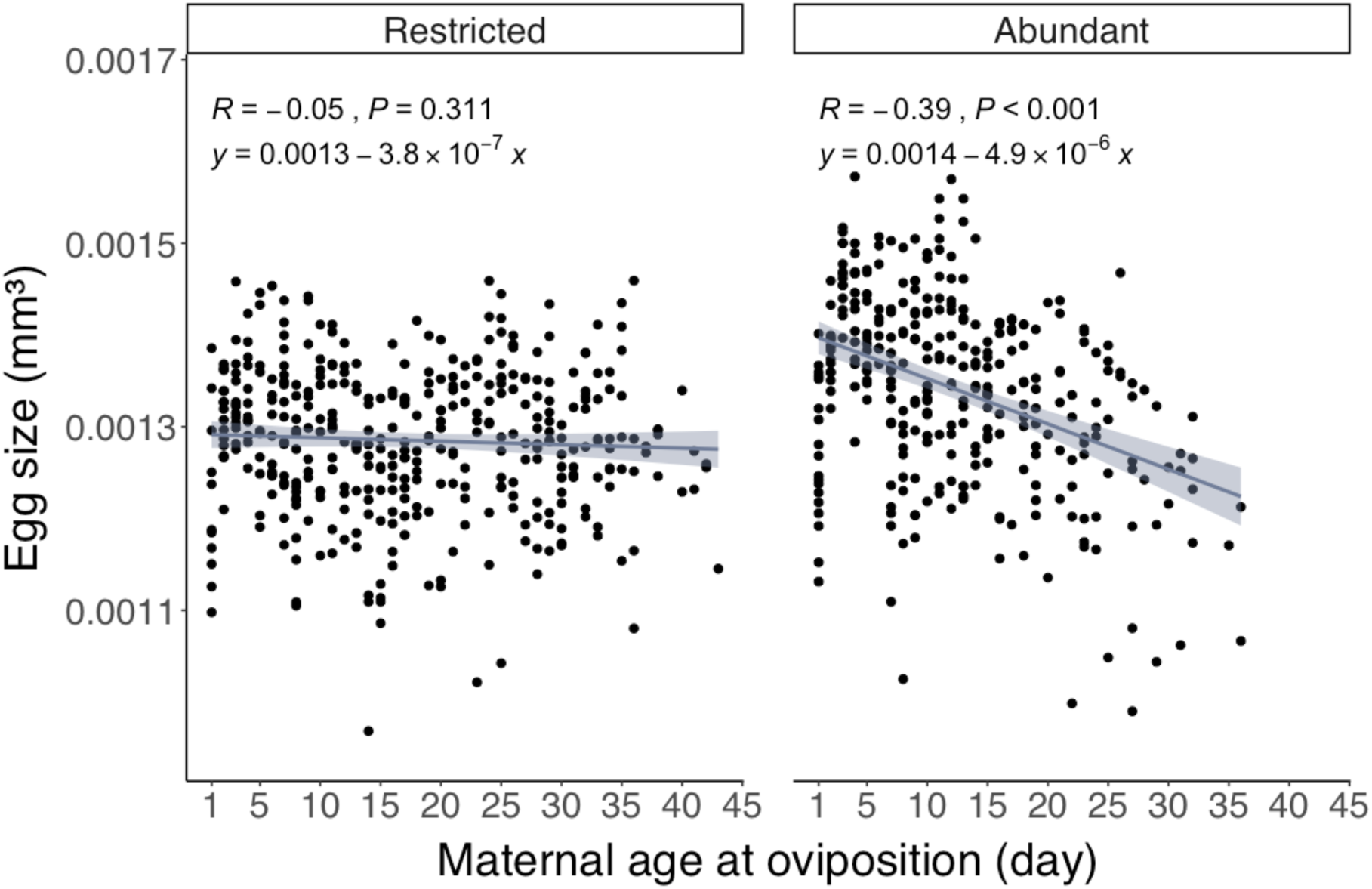
Egg size (volume) of *Amblyseius herbicolus* as influenced by maternal diet (restricted or abundant) and age at oviposition. Solid lines indicate regression fits, with shaded areas showing 95% confidence intervals. Regression equation, Pearson’s correlation coefficient (*R*), and *P*-value are provided.

### 3.2 Maternal diet and age influences on offspring parameters

#### 3.2.1 Hatching and survival to adulthood

Offspring were classified by maternal age at oviposition based on mean oviposition periods (TABLE S1). In the restricted group, ‘young’ offspring were derived from eggs laid on days 1 to 18, and ‘old’ from day 19 onwards; in the abundant group, ‘young’ were from days 1 to 13, and ‘old’ from day 14 onwards.

Hatching success was unaffected by maternal diet (logistic regression: Wald χ^2^ = 0.001, *df* = 1, *P* = 0.972), maternal age at oviposition (Wald χ^2^ = 0.576, *df* = 1, *P* = 0.448), offspring diet (Wald χ^2^ = 0.269, *df* = 1, *P* = 0.604), or their interactions (FIGURE 3A). In contrast, survival to adulthood was influenced by offspring diet (Wald χ^2^ = 83.683, *df* = 1, *P* < 0.001) and maternal age (Wald χ^2^ = 7.040, *df* = 1, *P* = 0.008), with a significant interaction between maternal age and offspring diet (Wald χ^2^ = 9.463, *df* = 1, *P* = 0.002). Specifically, offspring from older mothers and those provided more prey had higher survival, particularly under low prey availability (FIGURE 3B).

**FIGURE 3.**
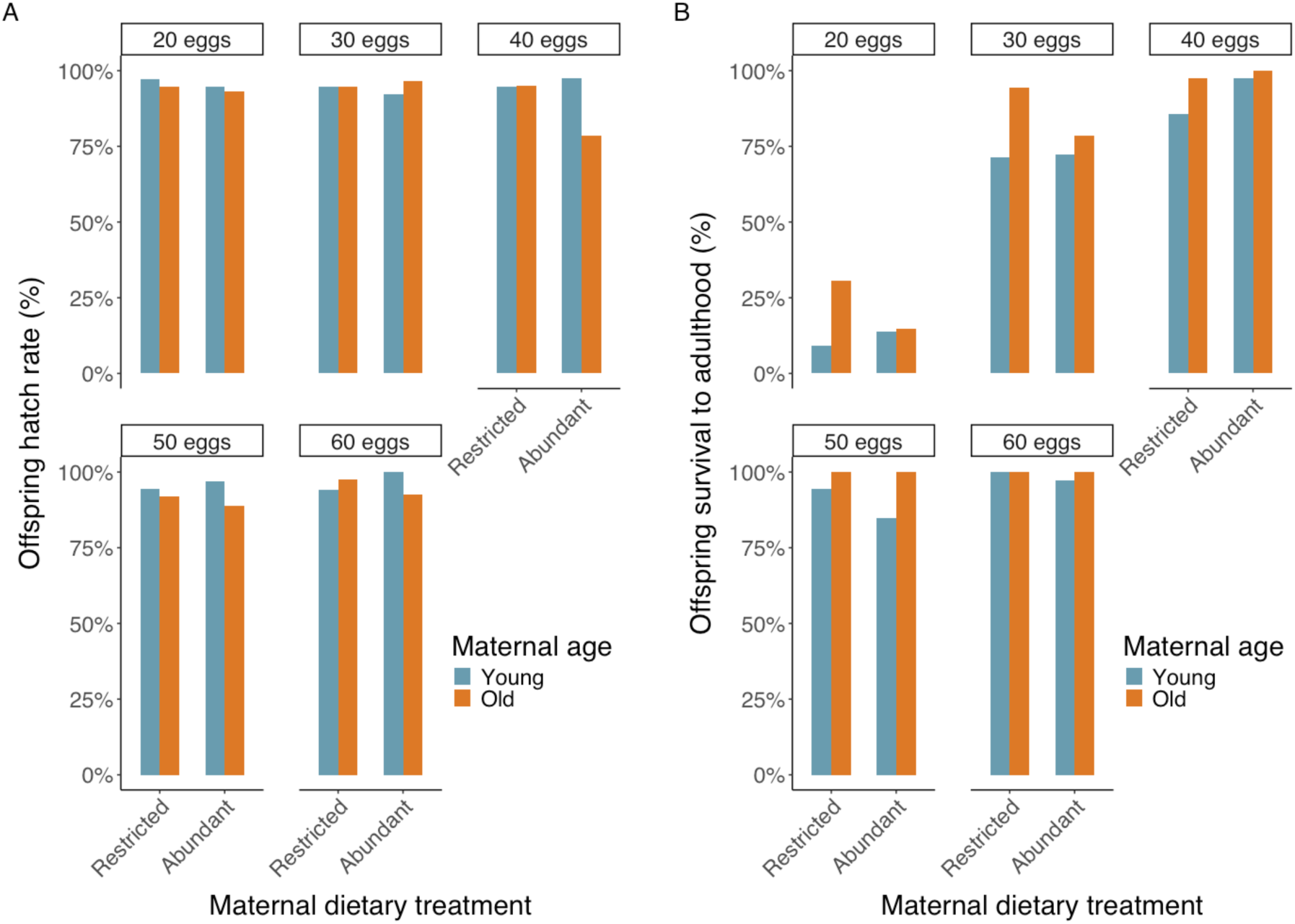
Hatching rate (A) and survival to adulthood (B) of *Amblyseius herbicolus* offspring under different offspring prey diets (20–60 eggs) and maternal age/diet treatments. Panels separate restricted and abundant maternal diets. Offspring from young and old mothers are defined as in the text.

#### 3.2.2 Developmental duration and size at maturity

Developmental duration was significantly affected by maternal age (LMM: Wald χ^2^ = 38.063, *df* = 1, *P* < 0.001) with a significant interaction between maternal age and diet (Wald χ^2^ = 21.710, *df* = 1, *P* < 0.001), but not maternal diet (Wald χ^2^ = 0.878, *df* = 1, *P* = 0.349) and prey consumption (Wald χ^2^ = 1.122, *df* = 1, *P* = 0.290). Offspring from older mothers took longer to mature, particularly in the abundant diet group (FIGURE 4).

**FIGURE 4.**
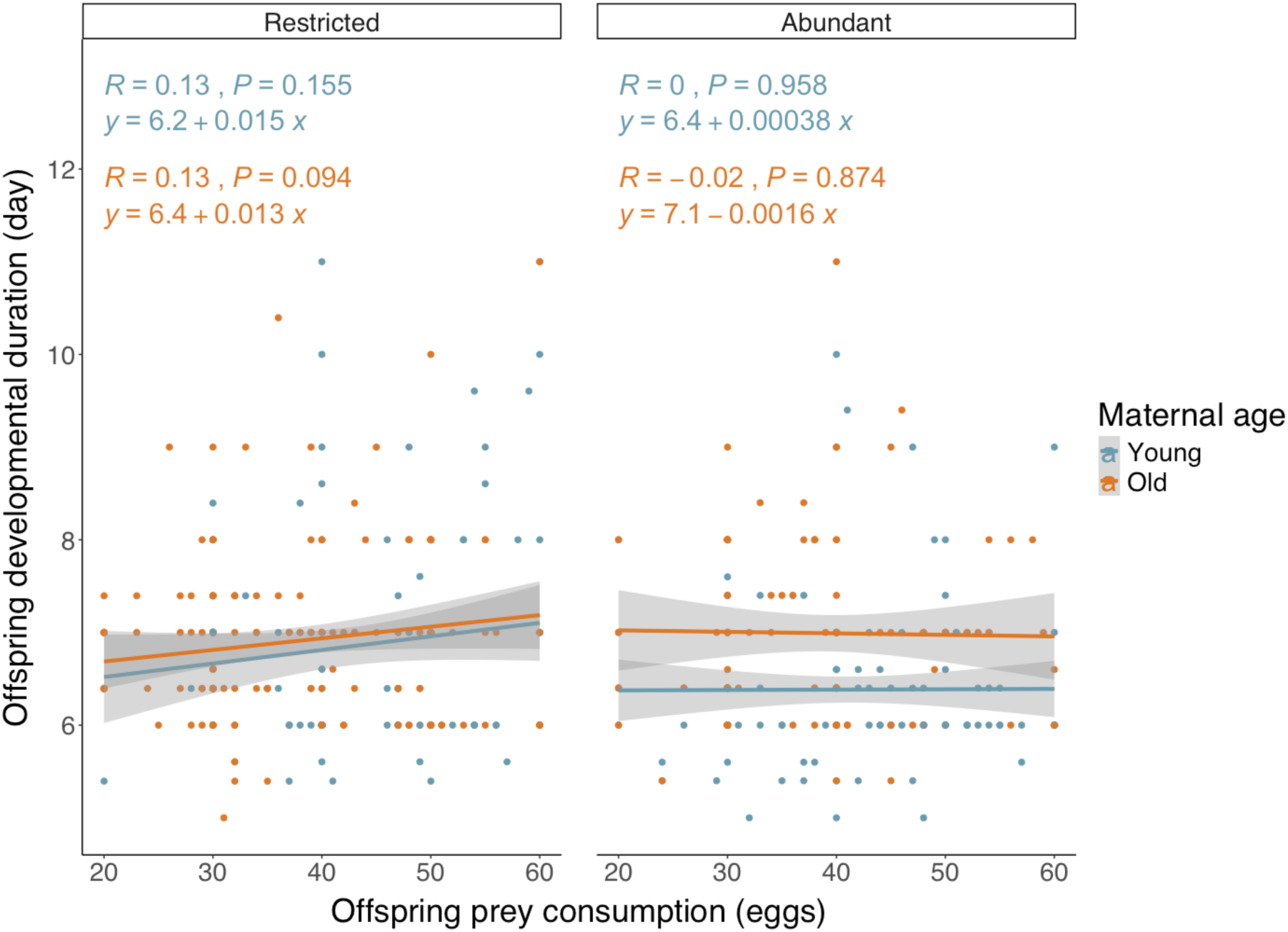
Developmental duration of offspring in relation to prey consumption, maternal diet, and maternal age at oviposition. Solid lines show regression fits; shaded areas indicate 95% confidence intervals. Offspring from young and old mothers are defined as in the text.

Size at maturity was significantly influenced by prey consumption (LMM: Wald χ^2^ = 56.224, *df* = 1, *P* < 0.001), maternal age (Wald χ^2^ = 4.238, *df* = 1, *P* = 0.040), and the interaction between maternal diet and age (Wald χ^2^ = 4.167, *df* = 1, *P* = 0.041). Larger size was associated with higher prey consumption and older mothers; differences between offspring of young and old mothers were more pronounced in the restricted maternal diet group (FIGURE 5).

**FIGURE 5.**
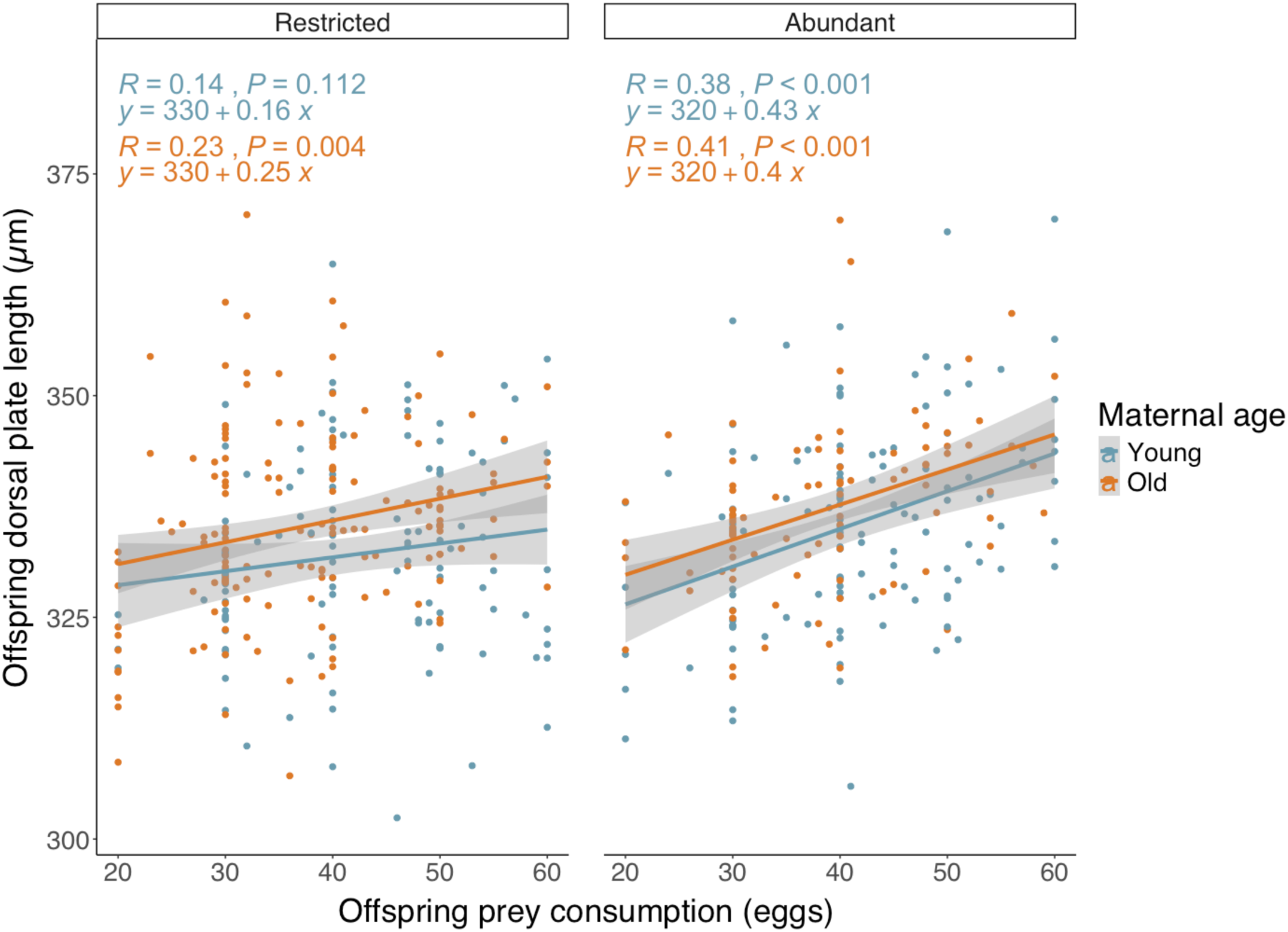
Dorsal plate length of offspring in relation to prey consumption, maternal diet, and maternal diet. Solid lines show regression fits; shaded areas indicate 95% confidence intervals. Offspring from young and old mothers are defined as in the text.

#### 3.2.3 Prey consumption

Offspring prey consumption increased with prey availability (GLM: Wald χ^2^ = 859.229, *df* = 1, *P* < 0.001). Maternal age at oviposition (Wald χ^2^ = 30.510, *df* = 1, *P* < 0.001) and the interaction between maternal diet and age (Wald χ^2^ = 4.288, *df* = 1, *P* = 0.038) were also significant: offspring of older mothers consumed less, especially when mothers were on the restricted diet (FIGURE 6). Maternal diet alone had no significant effect (Wald χ^2^ = 1.401, *df* = 1, *P* = 0.236).

**FIGURE 6.**
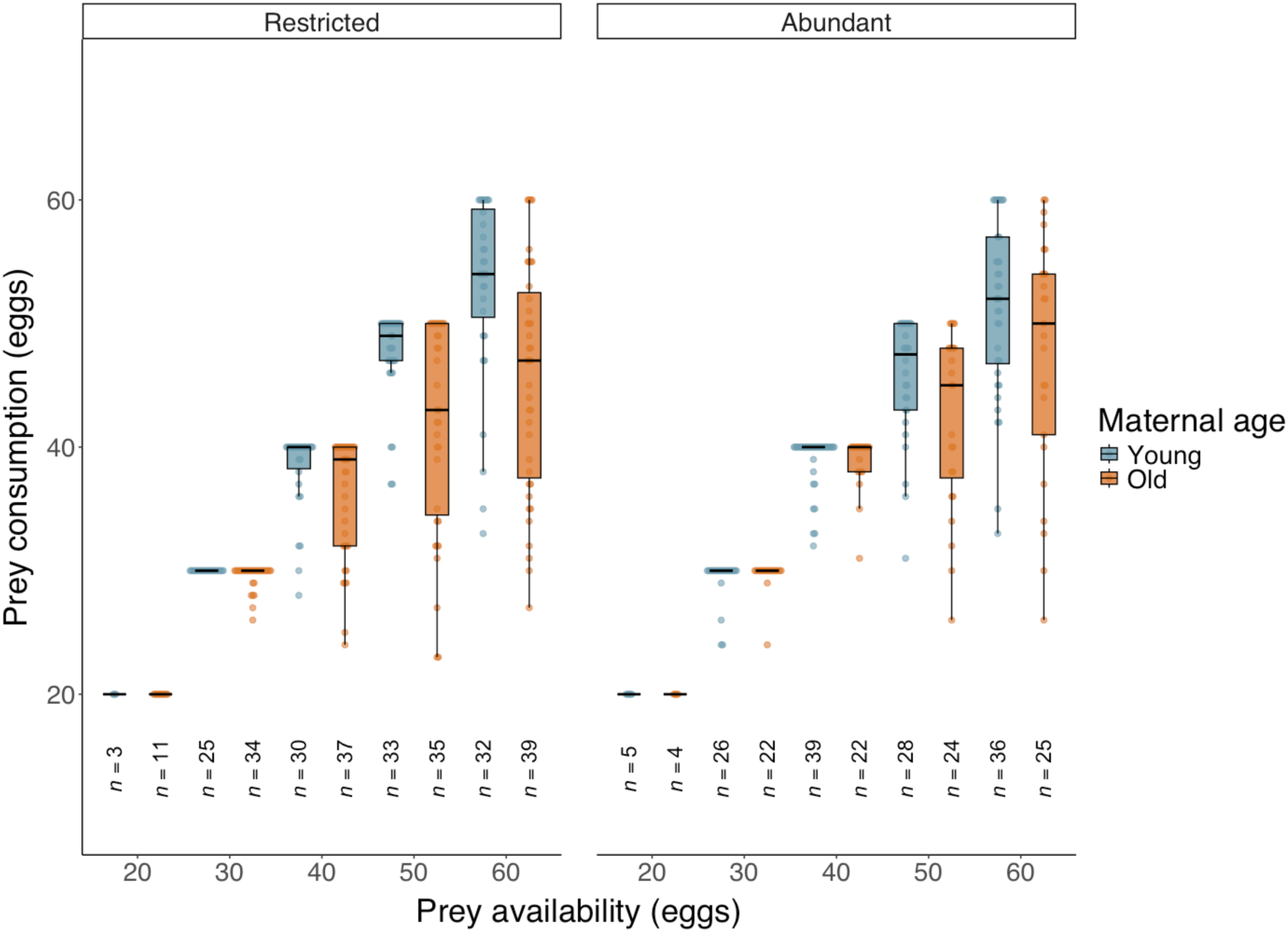
Offspring prey consumption by prey availability (20, 30, 40, 50, and 60 eggs), maternal diet, and maternal age. Individual points show observations (beeswarm); boxplots indicate median, interquartile range (IQR), and 1.5 × IQR whiskers. Panels separate restricted and abundant maternal diets. Sample sizes (*n*) are indicated below each box. Offspring from young and old mothers are defined as in the text.

## 4 Discussion

Our study provides evidence for an inverse Lansing effect in the predatory mite *A. herbicolus*. Offspring of older mothers showed higher survival to maturity, longer development, larger body size, and reduced prey consumption during development. Moreover, offspring of older mothers exhibited no significant reduction in developmental plasticity compared to those of younger mothers. These findings contrast with the classical Lansing effect and our first hypothesis, but align with studies showing that late-life reproduction can result in enhanced offspring quality in some species (Marshall et al., 2010; Travers et al., 2021; Anderson et al., 2022). For instance, older spiders (*Argiope radon*) produce offspring with greater starvation tolerance (Ameri et al., 2019), and beetles (*Menochilus sexmaculatus*) develop faster when produced by older mothers (Singh et al., 2021). Such findings suggest that maternal age effects may represent adaptive shifts in reproductive strategy rather than simple deterioration. Our results indicate a terminal investment strategy rather than senescent decline in *A. herbicolus*.

### 4.1 Maternal investment and offspring performance

Maternal diet strongly influenced life-history traits in *A. herbicolus*. Females on a restricted diet produced fewer eggs per day but extended their oviposition period and lived longer, resulting in lifetime fecundity comparable to females on an abundant diet. This trade-off between current reproduction and survival is consistent with resource-allocation models observed across taxa (Kaitala, 1991; Zera & Harshman, 2001; Lee et al., 2020).

Egg size and number were affected by both maternal diet and age. Larger eggs were produced under the abundant diet, which is consistent with other predatory mite species (Walzer & Schausberger, 2015). However, the reduction in egg size and number with advanced maternal age was only observed in those given the abundant diet. Such age-related patterns vary across taxa, with decreases in egg size in *Drosophila melanogaster* (Bloch Qazi et al., 2017) but increases in ladybirds (*Coleomegilla maculata*) (Vargas et al., 2012). Interestingly, diet-restricted females showed more stable investment across their reproductive lifespan, and we found no evidence for a size–number trade-off as observed in other species (Gliwicz & Guisande, 1992).

Despite reductions in egg size with maternal age, offspring performance was not compromised. Larger eggs are typically associated with enhanced offspring growth, size, and survival (Fox, 1994; Dias & Marshall, 2010; Rius et al., 2010). Thus, our finding suggests that maternal age effects in *A. herbicolus* operate independently of egg size. Similar findings have been observed in the soil mite *S. berlesei* (Benton et al., 2008). Since egg provisioning is an important mechanism underlying maternal age effects (Mousseau & Dingle, 1991; Mousseau & Fox, 1998; Yanchula & Alto, 2021), egg nutrient content and composition, rather than size alone, may be key factors in *A. herbicolus*. For example, older mothers of the parasitic wasp *Eupelmus vuilleti* showed reduced egg provisioning with protein, sugar, and lipids, which resulted in a lower nutrient composition at maturity (Muller et al., 2017). Testing whether maternal age alters egg nutrient composition in *A. herbicolus* remains an important future direction for research.

### 4.2 Maternal diet-by-age interactions

Maternal diet modulated the age effects on offspring. Differences in survival, size, and prey consumption between offspring of older and younger mothers were most pronounced when mothers had restricted diets. Contrary to our second hypothesis and previous models (Vijendravarma et al., 2010; Hafer et al., 2011; van den Heuvel et al., 2016), offspring from older, diet-restricted mothers had the best overall performance. One possible explanation is that the observed effects are anticipatory, where mothers prime their offspring against resource limitation. Alternatively, it could be that our restricted diet treatment was not sufficiently stressful to elicit phenotypic costs on offspring, and the mild dietary restriction may have buffered negative maternal age effects (Gribble et al., 2014; Hibshman et al., 2016). Stronger dietary stress could reverse this pattern, which should be investigated in future experiments.

The results of our study also highlight the context-dependence of maternal age effects, consistent with evidence that outcomes can vary across environmental conditions and taxa (Beckerman et al., 2006; Marshall et al., 2010; Kuijper & Johnstone, 2018; Zirbel & Alto, 2018).

### 4.3 Proximate mechanisms and ecological implications

The most important finding was that offspring of older mothers grew larger while consuming fewer prey, which suggests an increased efficiency of energy conversion (Abrams et al., 1996; Arendt, 1997). Whether this reflects altered metabolism, reduced energy expenditure (e.g., foraging activity), or other physiological mechanisms warrants further study.

Another possible explanation for reduced prey consumption between offspring of old and young mothers could be the variation in their predation strategy. In arachnids, superfluous killing—killing more prey than consumed—has been documented in both mites and spiders (Metz et al., 1988; Maupin & Riechert, 2001; López-Mercadal et al., 2024). Offspring of younger mothers might engage more in superfluous killing whereas those of older mothers might not. Although we found no direct evidence of partially consumed prey eggs, this remains a hypothesis for future testing.

Reduced prey consumption by the later-produced offspring could provide adaptive benefits. Later-produced offspring may encounter depleted resources because of utilisation by mothers and earlier-produced offspring. Lower consumption requirements may reduce intraspecific competition among siblings and with mothers, which enhances both maternal and offspring survival. As a solitary predator, reduced food demand in the later-produced *A. herbicolus* could also facilitate dispersal and colonisation of new patches. Maternal age may thus act as a cue of environmental change, enabling offspring to anticipate resource scarcity (Vargas et al., 2012) and the onset of environmental stresses (Rossi et al., 2016). Furthermore, such age-related changes in maternal provisioning strategies may enhance long-term fitness by diversifying brood performance (Cameron et al., 2017).

### 4.4 Future directions

Our study was conducted in an asexual system, which avoids confounding paternal effects but limits generalisation to sexually reproducing taxa (Castonguay & Angers, 2012; Verhoeven & Preite, 2014). Moreover, prey consumption declined with maternal age in both diet treatment groups. This dynamic could alter effective resource availability (i.e., more leftovers from older individuals) and influence maternal provisioning, potentially contributing to the observed inverse Lansing effect. Future work on sexually reproducing species would help determine whether similar age-related patterns and maternal provisioning dynamics occur under biparental reproduction.

The persistence of maternal-age effects across generations remains uncertain. Some studies report only immediate maternal effects, while others document transgenerational persistence (Plaistow et al., 2015; Goos et al., 2019; Wylde et al., 2019). Assessing whether maternal-age effects extend beyond a single generation, and whether these effects are additive or interactive with subsequent generations and environmental factors, can provide important insights into non-genetic inheritance.

Lastly, offspring may respond differently to variation in age-related maternal provisioning (Plaistow et al., 2015). Whether the observed inverse Lansing effect in *A. herbicolus* is a compensatory strategy, and whether this leads to trade-offs in later life, should be examined further.

## 5 Conclusions

In summary, we observed evidence of an inverse Lansing effect, where offspring of older mothers performed better than those of younger mothers during development. Our study highlights that maternal diet can affect offspring development in *A. herbicolus*, and that the effect depends strongly on maternal age at oviposition. The interaction between maternal diet and age suggests that maternal reproductive decisions and maternal effects are not static but change over time, and that the timing of oviposition plays a crucial role in shaping offspring outcomes. Maternal age may therefore play a key role in maintaining variation in life histories and should be considered more explicitly when examining maternal effects and offspring performance.

