## Supplemental Tables and Figures for "An inverse Lansing effect: Older mothers produce offspring with improved development and growth efficiency"

**Supporting information**

**TABLE S1.** Influence of dietary treatment (restricted vs. abundant frozen *Carpoglyphus lactis* eggs) on life-history traits of *Amblyseius herbicolus*. Restricted individuals received 30 prey eggs during immature development and 30 eggs replenished daily in adulthood, whereas abundant individuals received 50 prey eggs during immature development and 50 eggs replenished daily in adulthood. Values are presented as means ± SEMs. Asterisks indicate significant differences between dietary treatments from generalized linear models (GLMs): $P < 0.05$*, $P < 0.01$**, $P < 0.001$***.

| Measurement type | Parameter | Restriction (*n* = 17) | Abundant (*n* = 18) |
|---|---|---|---|
| Prey consumption | Development (prey eggs) | 28.8 ± 0.6*** | 44.2 ± 1.2*** |
| | Adulthood (prey eggs) | 759.1 ± 29.5* | 891.2 ± 47.0* |
| | Adulthood daily (prey eggs/day) | 18.6 ± 0.2*** | 29.9 ± 0.9*** |
| Durations | Developmental time (days) | 6.9 ± 0.2*** | 6.1 ± 0.1*** |
| | Pre-oviposition period (days) | 3.0 ± 0.1 | 2.8 ± 0.1 |
| | Oviposition period (days) | 37.1 ± 1.5*** | 25.9 ± 1.7*** |
| | Post-oviposition period (days) | 1.6 ± 0.3 | 1.6 ± 0.5 |
| | Lifespan (days) | 47.8 ± 1.7*** | 36.2 ± 1.6*** |
| Oviposition | Fecundity (eggs) | 43.4 ± 2.0 | 41.1 ± 2.6 |
| | Daily oviposition (eggs/day) | 1.2 ± 0.1*** | 1.6 ± 0.0*** |
| | Max. daily oviposition (eggs/day) | 2.0 ± 0.0*** | 2.6 ± 0.1*** |
| Size | Dorsal plate length (µm) | 343.2 ± 4.2* | 355.1 ± 3.7* |

GLM test statistics: for developmental prey consumption, Wald $\chi^2 = 56.705$, $df = 1$, $P < 0.001$; for adult prey consumption, Wald $\chi^2 = 5.696$, $df = 1$, $P = 0.017$; for adulthood daily prey consumption, Wald $\chi^2 = 169.180$, $df = 1$, $P < 0.001$; for developmental time, Wald $\chi^2 = 15.223$, $df = 1$, $P < 0.001$; for pre-oviposition period, Wald $\chi^2 = 3.649$, $df = 1$, $P = 0.056$; for oviposition period, Wald $\chi^2 = 22.197$, $df = 1$, $P < 0.001$; for post-oviposition period, Wald $\chi^2 = 0.003$, $df = 1$, $P = 0.955$; for lifespan, Wald $\chi^2 = 24.133$, $df = 1$, $P < 0.001$; for fecundity, Wald $\chi^2 = 0.496$, $df = 1$, $P = 0.481$; for daily oviposition, Wald $\chi^2 = 36.026$, $df = 1$, $P < 0.001$; for maximum daily oviposition, Wald $\chi^2 = 28.574$, $df = 1$, $P < 0.001$; for size, Wald $\chi^2 = 4.563$, $df = 1$, $P = 0.033$.

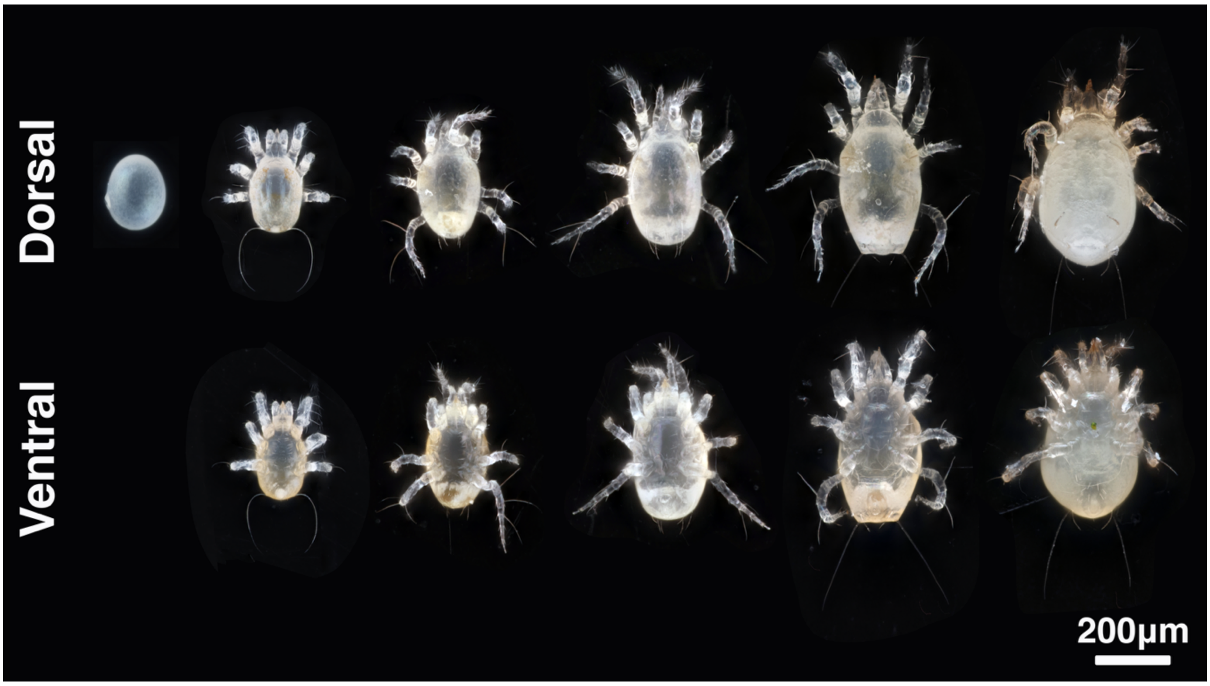


**FIGURE S1.** Dorsal and ventral views of all life stages (and phases) of *Amblyseius herbicolus*: egg, larva, protonymph, deutonymph, newly moulted adult female, and gravid adult female (left to right). Images captured with a 20× adapted lens.

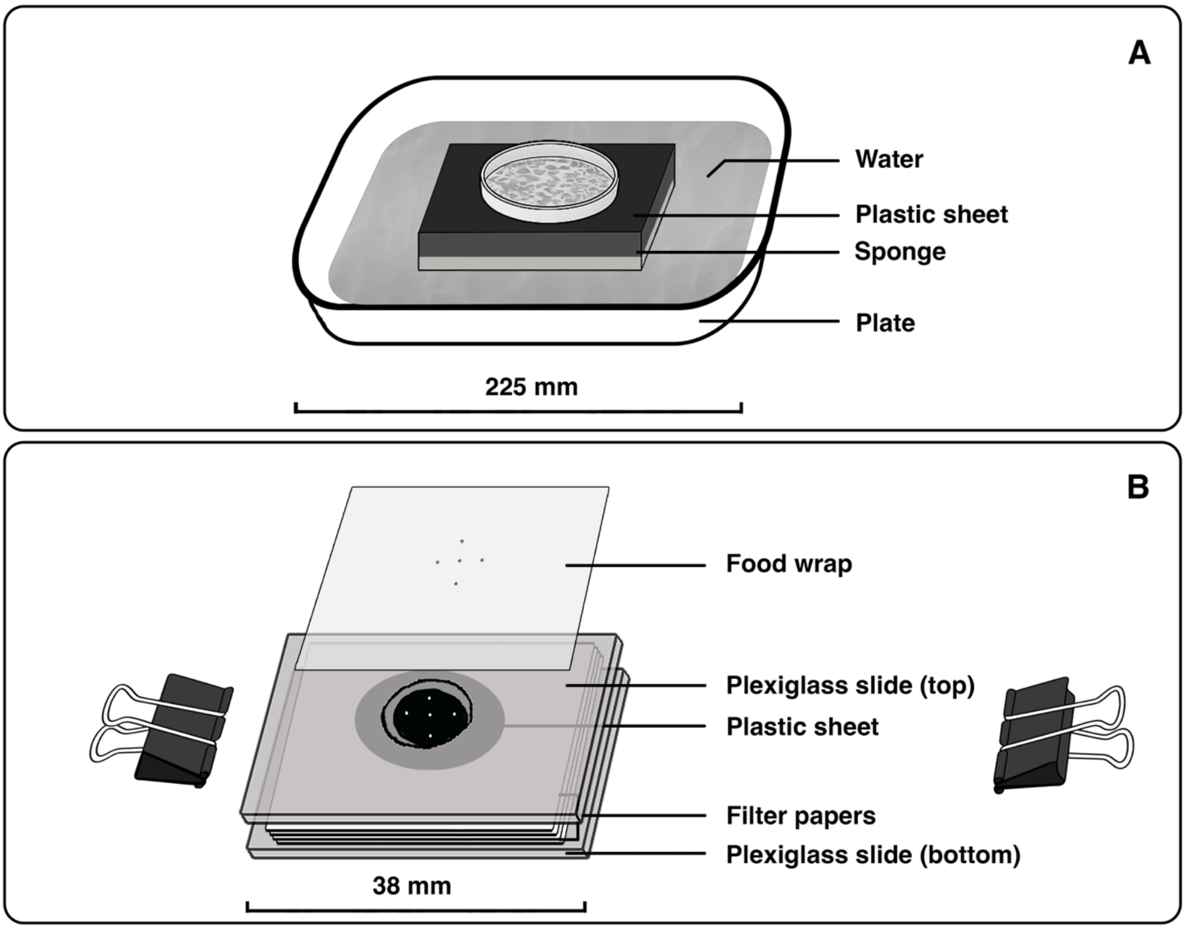


**FIGURE S2.** Rearing set-up and experimental units for *Amblyseius herbicolus*. A: Colony rearing container (225 × 225 × 35 mm, length × width × thickness) filled with water; a black plastic sheet on a sponge (100 × 100 × 30 mm) improved visibility and prevented overflow. *Carpoglyphus lactis* was reared in 55 mm Petri dishes containing wheat bran (~96%), sugar (~1%), and yeast (~3%). Folded black plastic sheets (~20 × 10 mm, length × width) and black threads (~20 mm long) provided resting sites for mites. B: Individual rearing cell composed of two plexiglass slides (38 × 25 × 2 mm, length × width × thickness). The top slide contained a cone-shaped hole (top diameter 6 mm, bottom 3 mm, volume 32.99 $mm^3$) covered with food wrap and black plastic film; small holes allowed air and moisture exchange. Filter paper beneath served as a water reservoir. Slides were held together with foldback clips.

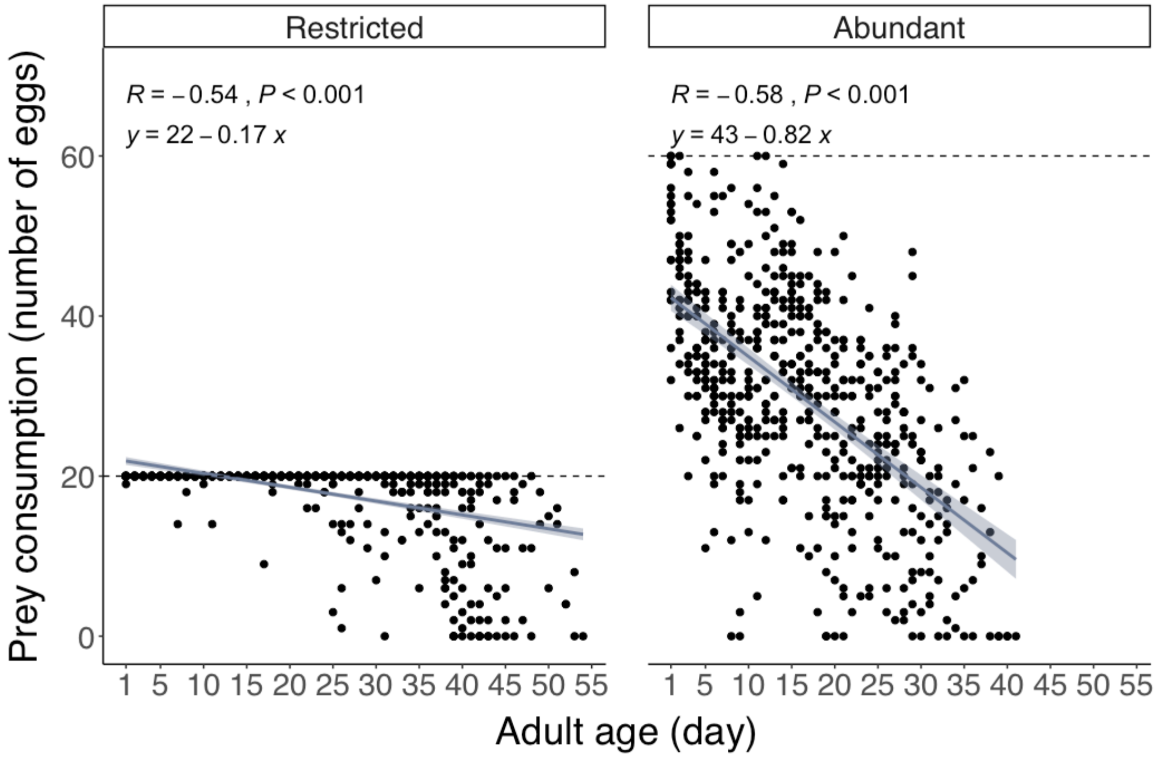


**FIGURE S3.** Effects of dietary regime (restricted vs. abundant *Carpoglyphus lactis* eggs) and maternal age on daily prey consumption of *Amblyseius herbicolus*. Dashed lines indicate the number of eggs provided per day (30 for restricted, 50 for abundant), replenished daily. Darker blue regression lines represent fitted trends with 95% confidence intervals (lighter blue). Regression equation, Spearman's correlation coefficient (*R*), and *P*-value are shown. Wider variation in the restricted group reflects a longer lifespan compared with the abundant group.

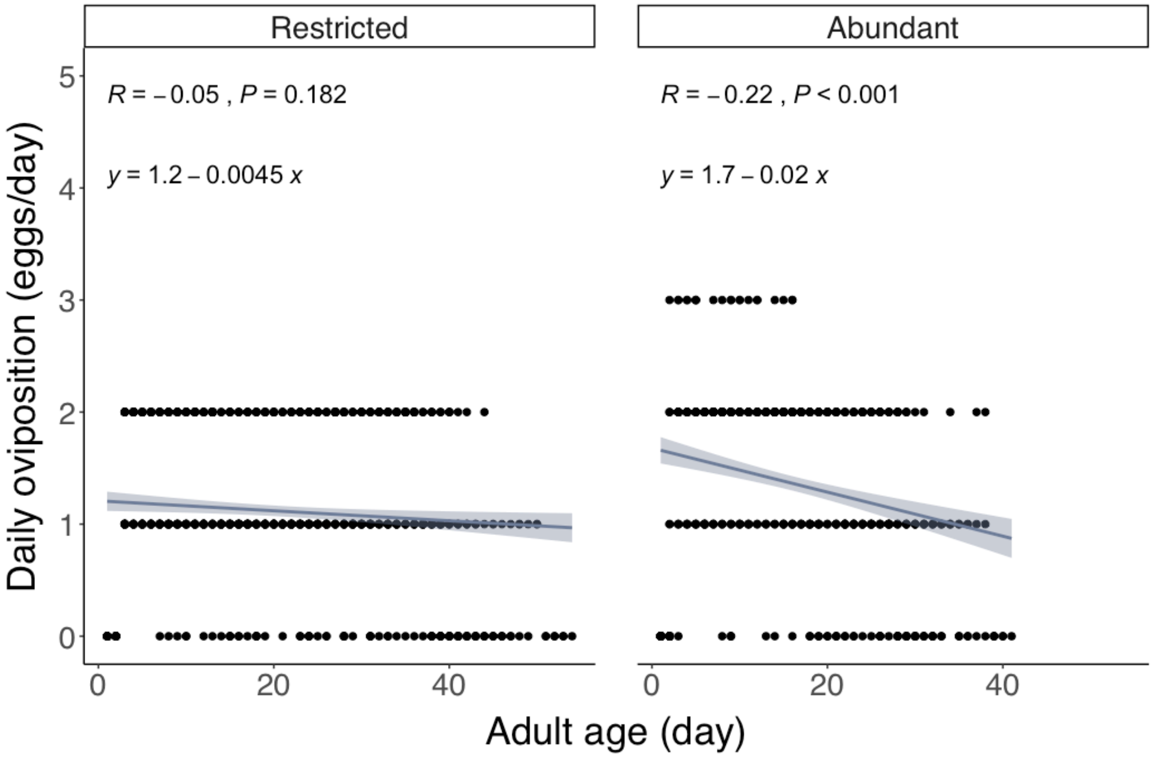


**FIGURE S4.** Effects of dietary regime (restricted vs. abundant *Carpoglyphus lactis* eggs) and maternal age on daily oviposition of *Amblyseius herbicolus*. Dashed lines indicate the number of eggs provided per day (30 for restricted, 50 for abundant), replenished daily. Darker blue regression lines represent fitted trends with 95% confidence intervals (lighter blue). Regression equation, Spearman's correlation coefficient (*R*), and *P*-value are shown. Wider variation in the restricted group reflects a longer oviposition period compared with the abundant group.
